# Protein Dimension DB: A Unified Protein Repository for Representation Learning and Functional Analysis

**DOI:** 10.1101/2025.09.29.679219

**Authors:** Pitágoras de Azevedo Alves Sobrinho, Tetsu Sakamoto, Wilfredo Blanco Figuerola

## Abstract

Inspired by the success of large language models in areas like natural language processing, researchers have applied similar architectures, notably the Transformer, to protein sequences. Thanks to these developments, Protein Language Models (PLMs) have become important resources for diverse tasks such as predicting protein family, function, solubility, cellular location, molecular interactions and remote homology. However, the size of the best performing PLMs (which can be up to 15B parameters) requires substantial computational power. Protein Dimension DB addresses this critical bottleneck by providing a centralized, version-controlled resource of precomputed protein embeddings, experimentally validated molecular function annotations, and taxonomic encodings. The database integrates embeddings from seven state-of-the-art PLMs, including ProtT5, ESM2, and Ankh variants for all Swiss-Prot/ UniProt proteins. These models were compared by benchmarking molecular function prediction. Tests revealed that hybrid embeddings (e.g., Ankh Base + ProtT5) outperformed single-model approaches with minimal dimensionality increases. Taxonomic encodings further boosted performance by 2.9% AUPRC, demonstrating lineage-aware learning. By providing embeddings in Parquet format — a columnar storage optimized for machine learning workflows — the resource eliminates GPU-dependent preprocessing and reduces storage requirements. This enables immediate use in resource-constrained environments while maintaining backward compatibility through versioned releases. All datasets are freely accessible via Github and HuggingFace, with unified metadata enabling applications from functional annotation to evolutionary studies. Protein Dimension DB bridges the gap between cutting-edge PLMs and practical biological research, offering researchers standardized inputs for reproducible, multi-modal protein analysis.

## 1 Introduction

The representation and annotation of proteins presents one of the most fundamental challenges in modern bioinformatics, with far-reaching implications in biomedical research, evolutionary biology, and biotechnology. Although recent breakthroughs in protein structure prediction, particularly AlphaFold2 [1], have revolutionized our understanding of protein folding, these methods remain computationally prohibitive for many research groups, requiring specialized hardware and days of processing time for comprehensive analyses [2]. This computational bottleneck is particularly acute for studies with a large amount of novel proteins, where the need to process thousands of protein sequences makes structure-based approaches impractical.

Protein Language Models (PLMs) have emerged as a powerful alternative, offering the ability to capture structural and functional information directly from amino acid sequences[3]. Models like ProtT5 [4] and ESM-2 [5] can generate informative protein representations with just seconds of computation on standard hardware [7]. These numerical representations (embeddings) encode evolutionary patterns, physicochemical properties, and potential functional motifs, providing a rich foundation for downstream predictive tasks [6]. This efficiency makes them particularly valuable for large-scale comparative genomics studies, rapid annotation of newly sequenced proteins, analysis of organisms with limited structural data and resource-constrained research environments.

The UniProtKB [8] project generates a dataset with ProtT5 embeddings for all proteins in their “Swiss-Prot” subset. However, there is a lack of per-protein embedding datasets using other PLMs. Researchers must typically generate embeddings anew for each study, wasting computational resources and introducing unnecessary variability. This creates unnecessary barriers to their widespread adoption in biological research.

The most recent Critical Assessment of Protein Function Annotation (CAFA) adopted NCBI taxonomic codes as one of its standard inputs, alongside protein sequences. Since then, several machine learning projects have used taxonomic unit encodings along with protein embeddings or structures [3,13], usually as “one-hot” encodings. Similarly to PLM embeddings, there is a lack of standardized taxon encodings available to the community.

This work has 3 main objectives:

‐ Enable researchers without access to high-end computing to use the encodings produced by the latest PLMs in their projects;
‐ To compare the capabilities of different PLMs;
‐ Measure how much strategies such as the combination of different PLMs and taxonomic information can improve molecular function prediction;

We approached these challenges with the creation of Protein Dimension DB, a resource which includes several datasets for the proteins in Swiss-Prot/Uniprot, such as PLM-based embeddings, numerical representations of taxonomy and experimentally confirmed gene ontology annotations.

The datasets in Protein Dimension DB were used to create molecular function prediction models, which were benchmarked. The Ankh family of models showed the highest performance. The tests also revealed that pairing medium-sized PLMs can provide the same level of performance as using the largest models and that taxonomic information can improve performance by up to 2%.

## 2 Building The Database

The data pipeline was implemented as a Nextflow pipeline. Data collection starts with obtaining the complete Swiss-Prot sequences dataset, which comprises the highest quality protein sequences of UniprotKB [8]. It accounted for a total of 569,059 proteins. The Gene Ontology Annotation (GOA) Database was downloaded and then filtered to exclude electronic annotations (IEA) and only keep annotations of Swiss-Prot proteins [10].

ProtT5 protein embeddings are downloaded from the UniprotKB database and converted from the original HDF5 format to Parquet format. Parquet format was used because its columnar storage design, which enables efficient compression and faster column-wise queries compared to row-based formats like HDF5 [11]. This is particularly beneficial for machine learning workflows, where selective access to specific columns and rows is common. This format was also adopted for the remaining datasets.

Differently from ProtT5 - Ankh and ESM2 embeddings were not available for all Swiss-Prot proteins. To address this, they were calculated locally. Ankh had two variants available: Base and Large. The ESM2 project had several different models: ESM2 T6, T12, T30, T33 and T36. Both PLM projects have Python APIs [12,5], which we used to transform protein sequences into embeddings. The smaller models (ESM2 T6, T12, T30 and Ankh Base) showed good performance, but inference on the larger models (ESM2 T36 and Ankh Large) required several weeks of processing.

The NCBI taxon ID of each protein was also obtained from UniprotKB. The frequency of each taxa was counted and two sets were selected: The 128 and 256 most common taxa. They were treated as discrete categories and converted to “onehot” encodings.

## 3 Case Study: Benchmarking of Protein Molecular Function Classification

To evaluate the different protein representations and demonstrate the utility of Protein Dimension DB, we conducted a comprehensive benchmark evaluating different combinations of protein embeddings and taxonomic encodings for molecular function (MF) prediction. Using the experimentally validated GO annotations from GOA as ground truth, we trained a multi-label classifiers to predict MF terms, measuring performance via weighted AUPRC (prioritizing rare classes) and ROC AUC.

### 3.1 Protein and Molecular Function Sampling

First, we filtered the full list MF terms, keeping only those annotated at least 36 different proteins. This resulted in 1,487 MF terms. A total of 102,696 proteins (18% of Swiss-Prot) were annotated to at least one of those terms. Of these, 15% (15,577) was randomly selected for validation, 30% (30,808) for testing and 55% (56,311) for training.

We wanted to measure how these models perform not only on functions with a vast number of protein annotations, but also on functions with very few. Following this principle, we sorted the 1,487 trainable MF terms by their frequency. Then the first, middle and top 24 terms were selected. Combined, they made up a list of 72 MFs, which were used as classification targets.

### 3.2 Model Architecture

The classifier is a deep neural network (DNN). It was based on the architecture of the PROTGOAT model [3], due to its modular approach (Figure 1). The model features separate modules for the different input features, whose last layers are connected by a concatenation layer, followed by a final Relu-activated dense layer and a Sigmoid-activated layer as output. Training used Adam optimizer and Binary Cross-Entropy as loss function. This design can be used to make classifiers with any list of input features, when properly optimized.

**Fig. 1.**
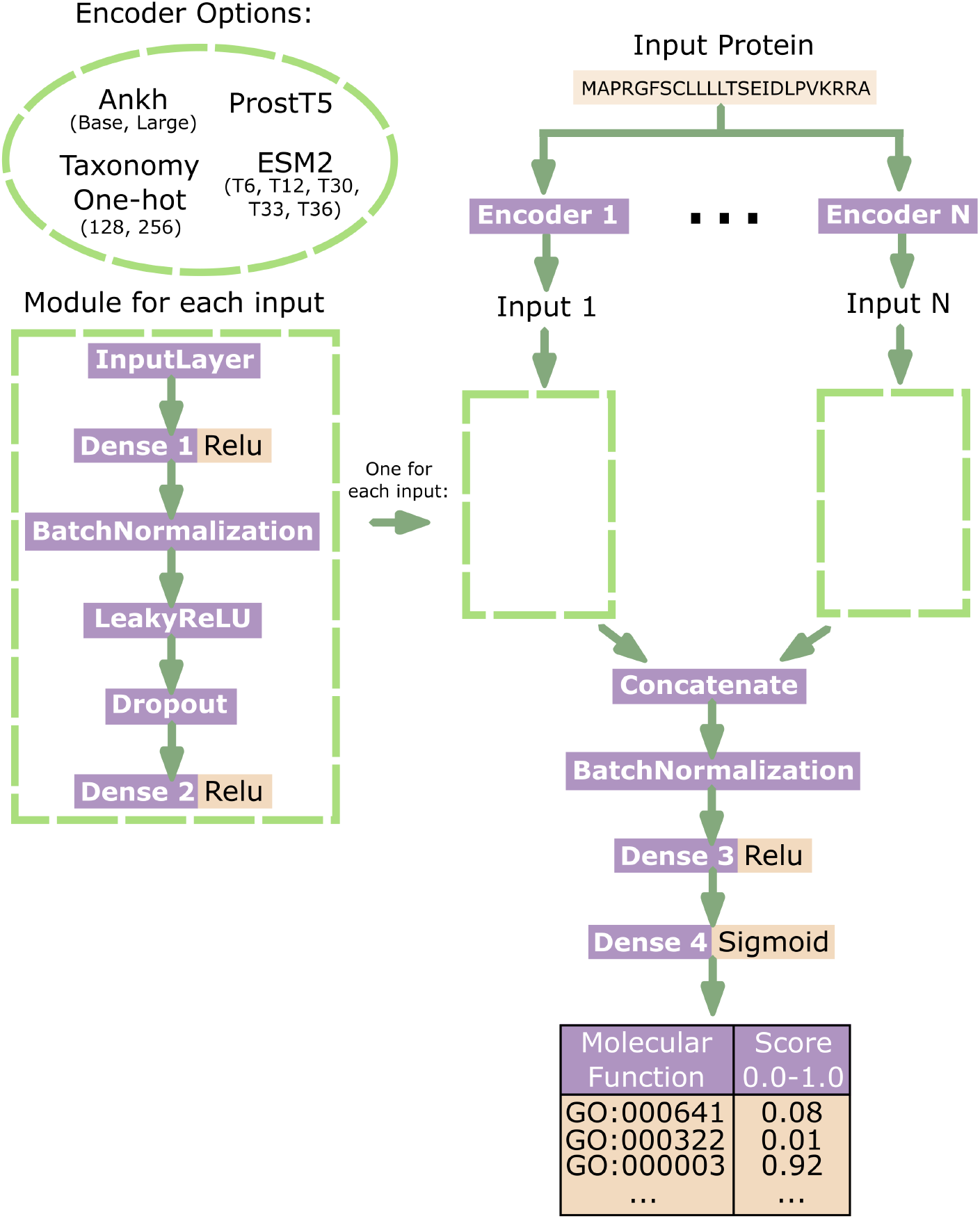
Our implementation of the DNN architecture proposed by PROTGOAT [3]. The modular neural network architecture processes input features (such as protein embeddings and taxonomic encodings) through separate feature-specific branches. Each input type passes through two dense layers with batch normalization, LeakyReLU activation and dropout regularization. Intermediate representations are concatenated then refined through final dense layers (ReLU-activated) before sigmoid-activated output for multi-label molecular function prediction. This design enables flexible integration of heterogeneous biological data while mitigating overfitting through pathway-specific regularization.

### 3.3 Optimization and Benchmarking

Metaparameter optimization was performed with random search over a uniform distribution, with each metaparameter having a minimum and maximum value (Table 1). Each feature had specific metaparameters (such as the dimensions of layers “Dense 1” and “Dense 2”), which were selected separatly.

**Table 1.** Metaparameter Ranges Used for Optimization.

| Metaparameter | Min. | Max. |
| --- | --- | --- |
| "Dense 1" width | 64 | 3600 |
| "Dense 2" width | 64 | 2048 |
| "Dense 3" width | 300 | 550 |
| LeakyRELU Negative Slope | 0.03 | 0.9 |
| Dropout Rate | 0.2 | 0.5 |
| Patience | 8 | 13 |
| Epochs | 38 | 48 |
| Learning Rate | 0.0006 | 0.00075 |
| Batch Size | 130 | 200 |

Each generation of the optimization consisted of 120 random settings, made using the default minimum and maximum values. The features of the training proteins were used to train one model for each metaparameter setting. Then, the fitness of each model was calculated on the test proteins, by comparing the generated scores with the ground truth labels. Two metrics were calculated: ROC AUC Score and AUPRC Score, weighted to account for class imbalance. The fitness was the average of the two metrics: (ROC AUC + AUPRC) / 2.

At the end of the generation, the 80 settings with best fitness were selected and the maximum and minimum range values of metaparameters was updated, using the minimum and maximum found in the best performing settings. This was done to gradually find better bounds metaparameter ranges for each feature.

After 4 generations, the setting with higher fitness was selected for validation. Validation was performed by predicting scores on the validation proteins, with weighted ROC AUC and AUPRC scores.

First, this metaheuristic algorithm was used to benchmark each individual PLM embedding as as the single input feature. Subsequent analysis paired the top five best-performing PLMs (Ankh Large, Ankh Base, ESM2 T36, ProtT5 and ESM2 T33) to identify optimal combinations of two different PLMs (Table 2). Combinations with lower performance PLMs were not tested due to hardware and time constraints. The three highest-performing pairs showed comparable results, with the Ankh Base + ProtT5 combination selected for further testing due to its more compact input dimensionality (1792 values total). This pairing was then augmented with onehot encodings of either 128 or 256 most frequent taxonomic groups as a third input feature.

**Table 2.** Molecular Function Classification Benchmark.

| Input Features Used | Total Feature Length | AUPRC Score (W) | ROC AUC Score (W) | F1 Score (W) |
| --- | --- | --- | --- | --- |
| TAXA 256 + Ankh Base + ProtT5 | 2048 | 0.8006 | 0.9094 | 0.7507 |
| TAXA 128 + Ankh Base + ProtT5 | 1920 | 0.796 | 0.9051 | 0.7487 |
| Ankh Base + ESM2 T33 | 2048 | 0.7802 | 0.8888 | 0.7283 |
| Ankh Base + Ankh Large | 2304 | 0.7796 | 0.8892 | 0.7317 |
| Ankh Base + ProtT5 | 1792 | 0.778 | 0.8905 | 0.7289 |
| Ankh Large + ProtT5 | 2560 | 0.7767 | 0.8913 | 0.7307 |
| Ankh Large + ESM2 T36 | 4096 | 0.7765 | 0.8889 | 0.7252 |
| Ankh Base + ESM2 T36 | 3328 | 0.776 | 0.8854 | 0.7266 |
| Ankh Large | 1536 | 0.7753 | 0.8889 | 0.7272 |
| ESM2 T36 + ProtT5 | 3584 | 0.7722 | 0.8884 | 0.7195 |
| Ankh Base | 768 | 0.7718 | 0.8831 | 0.7238 |
| Ankh Large + ESM2 T33 | 2816 | 0.7714 | 0.8851 | 0.7249 |
| ESM2 T33 + ProtT5 | 2304 | 0.7704 | 0.8883 | 0.72 |
| ESM2 T36 | 2560 | 0.7641 | 0.8824 | 0.7104 |
| ESM2 T33 + ESM2 T36 | 3840 | 0.7636 | 0.8811 | 0.7143 |
| ProtT5 | 1024 | 0.7604 | 0.8842 | 0.7142 |
| ESM2 T33 | 1280 | 0.7583 | 0.8817 | 0.7075 |
| ESM2 T30 | 640 | 0.7466 | 0.8765 | 0.6967 |
| ESM2 T12 | 480 | 0.739 | 0.8707 | 0.6861 |
| ESM2 T6 | 320 | 0.7162 | 0.8606 | 0.6638 |

The table presents Total Feature Length (sum of the lengths of all features used), Area Under the Precision-Recall Curve (AUPRC) Score, Area Under the Receiver Operating Characteristic Curve (ROC AUC) Score and F1 Score for all feature combinations tested. The AUPRC, ROC AUC and F1 metrics were calculated using the "weighted" averaging mode of the Scikit-Learn library [16], in order to address class imbalance.

Our benchmarking revealed several key insights. The top-performing configuration combined Ankh Base, ProtT5 embeddings, and 256-taxon encodings, achieving an AUPRC of 0.8006. This performance gain stems from Ankh’s and ProtT5’s protein embeddings combined performance, while taxonomic encodings provided an additional 2.9% AUPRC improvement over embedding-only models. Interestingly, the difference between using 128 versus 256 taxonomic groups proved minimal (<0.5% AUPRC), suggesting diminishing returns beyond the most prevalent taxa.

## 4 Discussion

Model size comparisons revealed important trade-offs. While larger ESM-2 variants (e.g., T36 with 2560 dimensions) marginally outperformed smaller ones (T6 with 320 dimensions), they required eight times the feature dimensionality. Notably, Ankh Large (1536 dimensions) surpassed ESM-2 T36 in AUPRC (0.7753 vs. 0.7641) despite its more compact representation. Model pairs showed higher performance when compared to single-model approaches. However, certain pairs like Ankh Large + ESM-2 T36 (4096 dimensions) demanded significantly more memory without delivering proportional performance gains.

The fact that the combination of embeddings from different PLMs (such as Ankh Base + ProtT5) outperform single models suggests that different models may capture complementary information about the protein that is relevant for predicting molecular function. Although it is difficult to specify exactly which features are described in the embedding space of each model, one can hypothesize that they learn different representations or prioritize different aspects of the sequence or structure that, when combined, provide a more complete view of the protein and its function.

For example, one model may be more effective in capturing local patterns important for binding sites or catalytic activity, while another may capture global or structural features important for subcellular localization or protein-protein interactions. ProtT5, for example, has been shown to capture aspects of protein structure impressively and to be effective in predicting variant effects [17].

The taxonomic encodings proved particularly valuable. According to Tiittanen et al.[15], taxonomic characteristics may improve classification because they allow models to generate different predictions for the same input sequence when the sequence occurs in different regions of the species taxonomy tree. However, Swiss-Prot contains a disproportional amount of sequences from *Homo sapiens* and model organisms. This represents an annotation bias, so results may not generalize equally well to proteins from underrepresented taxonomic groups.

## 5 Conclusions and Future Work

The results of the case study on molecular function prediction align with those of Vieira et al. [7], who concluded that medium-sized PLMs can achieve performance comparable to larger models while requiring substantially fewer computational resources. In addition, taxonomic input significantly improved function predictions.

These findings collectively demonstrate how precomputed features facilitate efficient experimentation with multi-modal inputs. By providing pre-generated embeddings from seven state-of-the-art PLMs, the database eliminates the need for researchers to perform resource-intensive computations, making advanced protein analysis accessible to groups without specialized hardware.

All molecular function annotations are filtered to exclude low-confidence predictions, ensuring that researchers work with reliable GO term assignments. This curation is particularly valuable given the noise present in many automated annotation pipelines [14].

Version-controlled releases in machine-learning-ready formats (Parquet) ensure consistency across studies, supporting the growing emphasis on reproducible research in computational biology. The datasets, which are currently in version 1.0, can be downloaded individually from the project page: https://pentalpha.github.io/protein_dimension_db/. By making these resources freely available, we aim to accelerate discoveries across diverse areas of protein science, from basic biological research to applied biotechnology and drug discovery.

As the field continues to develop new PLM architectures and annotation strategies, this resource will serve as a foundation for integrating these advances into practical research workflows, helping to realize the full potential of machine learning in protein science.

## Acknowledgments

We were able to generate the embeddings included in this database thanks to the computational infrastructure available in the Bioinformatics Multidisciplinary Environment (BioME), of the Federal University of Rio Grande do Norte.

